# The AFB1 auxin receptor controls rapid auxin signaling and root growth through membrane depolarization in *Arabidopsis thaliana*

**DOI:** 10.1101/2021.01.05.425399

**Authors:** Nelson BC Serre, Dominik Kralík, Ping Yun, Sergey Shabala, Zdeněk Slouka, Matyáš Fendrych

## Abstract

The existence of an electric gradient across membranes is essential for a cell operation. In plants, application of the growth regulator auxin (IAA) causes almost instantaneous membrane depolarization in various cell types, making membrane depolarization a hallmark of the rapid non-transcriptional responses to IAA. Auxin triggers rapid root growth inhibition; a process that underlies gravitropic bending. The growth and depolarization responses to auxin show remarkable similarities in dynamics, requirement of auxin influx and the involvement of the TIR1/AFB auxin coreceptors, but whether auxin-induced depolarization participates in root growth inhibition remains unanswered. Here, we established a toolbox to dynamically visualize membrane potential *in vivo* in *Arabidopsis thaliana* roots by combining the DISBAC2(3) fluorescent probe with microfluidics and vertical stage microscopy. This way we show that auxin-induced membrane depolarization tightly correlates with rapid root growth inhibition and that the cells of the transition zone/early elongation zone are the most responsive to auxin. Further, we demonstrate that auxin cycling in and out of the cells through AUX1 influx and PIN2 efflux is not essential for membrane depolarization and rapid root growth inhibition but acts as a facilitator of these responses. The rapid membrane depolarization by auxin instead strictly depends on the AFB1 auxin receptor, while the other TIR1/AFB paralogues contribute to this response. The lack of membrane depolarization in the afb1 mutant explains the lack of the immediate root growth inhibition. Finally, we show that AFB1 is required for the rapid depolarization and rapid growth inhibition of cells at the lower side of the gravistimulated root. These results are instrumental in understanding the physiological significance of membrane depolarization for the gravitropic response of the root and clarify the role of AFB1 as the receptor central for rapid auxin responses, adding another piece to the puzzle in understanding the biology of the phytohormone auxin.

## Introduction

Membrane potential reflects the difference between cytoplasmic and apoplastic electrical potentials and is conferred by operation of transporting proteins at the cellular membranes. The existence of a voltage gradient is essential for cellular operation and metabolism as this electrical difference allows for secondary transport processes across the membrane. In plants, the resting membrane potential is negative inside the cell (typically −120 to −160mv, Sze et al., 1999) and is fueled by the activity of the plasma membrane proton pumps (Sze et al., 1985). The application of the plant growth regulator auxin (IAA) causes almost instantaneous membrane depolarization in various cell types (Goring et al., 1979; Bates and Goldsmith 1983; Felle et al., 1991; Tretyn et al., 1991; Dindas et al., 2018, 2020; Paponov et al., 2019). This qualifies depolarization as a hallmark of the rapid non-transcriptional responses to IAA. Using the model of *Arabidopsis thaliana* root hairs, Dindas et al., (2018) demonstrated that the depolarization response requires functional IAA transport and signaling by the TIR1/AFB IAA co-receptors. The physiological significance of the auxin-triggered depolarization is however not understood. Specifically, in root cells, application of auxin triggers rapid root growth inhibition (RGI, Evans et al., 1994; Monshausen et al., 2011; Scheitz et al., 2013); a process that is required for the gravitropic bending. RGI requires intracellular auxin accumulation driven by the AUX1 influx carrier and is initiated by the TIR1/AFB coreceptors, AFB1 paralogue playing a crucial role (Fendrych et al., 2018; Prigge et al., 2020). The nature of the rapid TIR1/AFB(1) signaling branch is unknown, as well as the molecular machinery that executes the auxin-triggered growth inhibition. The growth and depolarization responses to auxin show remarkable similarities in timing, requirement of AUX1 and the enigmatic involvement of the TIR1/AFB coreceptors.

Here we established and optimized a non-invasive method to dynamically record membrane potential in plant roots, and we combined it with genetic and pharmacological approaches, as well as automated analysis of gravitropic response in *Arabidopsis thaliana* roots. This allowed us to answer the following longstanding questions. Is the transport of auxin across the membrane directly involved in membrane depolarization? What is the role of TIR1/AFB auxin receptors in membrane depolarization? And finally, what is the significance of membrane depolarization for root growth inhibition and gravitropism?

## Results and discussion

As the traditional membrane potential measurement by impaling cells with microelectrodes is incompatible with undisturbed root growth, we tested the membrane potential fluorescent dye DISBAC_2_(3) (Renier et al., 1995) to optimize simultaneous detection of growth and membrane potential in *Arabidopsis thaliana* roots. We imaged growing roots using a vertical-stage (von Wangenheim et al., 2017) spinning disc microscope. The voltage sensor-stained root cells, with a strong signal in the outer tissues epidermis and cortex and a nonspecific signal in the dead lateral root cap cells (Fig.S1a). DISBAC_2_(3) staining was not toxic as judged by the growth of the primary root (Fig.S1b,c). To test whether it reported changes in membrane potential, we quantified the effects of two known membrane polarization disruptors on the inner epidermis/outer cortex cells fluorescence. Application of fusicoccin (FC), an activator of the plasma membrane proton pump (Baunsgaard et al., 1998), decreased the DISBAC_2_(3) fluorescence indicating hyperpolarization (Fig.S1d,e; Senn and Goldsmith 1988; Ulrich and Novacky 1990). On the other hand, Carbonyl cyanide m-chlorophenyl hydrazone (CCCP), an uncoupler that dissipates the proton gradient (Labady Jr. et al., 2002), significantly increased the fluorescence reporting a depolarized state (Fig.S1d,e). Finally, to gain insight into the detection range of DISBAC_2_(3), we compared direct membrane potential measurements (using an impaled microprobe) with the fluorescence measurements in response to a KCl gradient (Fig.1a, S1f). Both methods recorded similar profiles. However, DISBAC_2_(3) displayed lower sensitivity in the detection of deep hyperpolarized states, with no detectable differences between 0.1mM and 1mM KCl. Nevertheless, the range of significant fluorescence detection was covering the fluorescence observed in the control, FC and CCCP treatments (Fig.1a). Taken together, these results indicated that DISBAC_2_(3) can report membrane potential variation in the physiological range of *Arabidopsis thaliana* root cells and can be used to non-invasively monitor spatiotemporal kinetics of membrane potential changes.

Auxin triggers growth inhibition within ∼30 seconds and this rapid and reversible response lasts for tens of minutes (Fendrych et al., 2018). Therefore, we measured the steady state growth and membrane potential response (>20 min after IAA application) by imaging seedlings in microscopy chambers containing agar medium with the respective treatments. To capture the initial phases of the response (<5 minutes), we imaged the seedlings in closable microfluidic chips (Fig.S1g,h). Applied at 100nM, the native auxin IAA triggered a spatially heterogeneous depolarization in *Arabidopsis thaliana* roots (Fig.1b, Sup. movie 1). Root hairs and epidermal cells in the transition/elongation zones showed strongest depolarization after 5 minutes of treatment. For this reason, we decided to focus on membrane potential responses in the cells of the transition zone for further experiments. Moreover, this zone is a hotspot of root growth regulation (Verbelen et al., 1996). To analyze whether membrane depolarization is proportional to RGI, we analyzed root response to a gradient of IAA auxin (0, 10, 100, 1000 nM; Fig.1c). In steady state, growth was strongly inhibited by 10nM IAA (Fig.1d), while a significant depolarization was only detected at high IAA concentrations (100 and 1000nM; Fig.1e). We further focused on the initial stages of the response. Application of 10 nM IAA (or higher) triggered an instantaneous (<30s) membrane depolarization and the response amplitude was positively correlated with IAA concentration (Fig.1f,g). In parallel, RGI began instantly (<30s) but the amplitude was similar between concentrations after 5 minutes (Fig.1h,i). Benzoic acid did not elicit membrane depolarization (Fig.1f,g) and did not inhibit growth (Fig.1h,i), apart from a partial inhibition at a high concentration (Fig.1d). These results show that physiological IAA levels elicit membrane depolarization and growth inhibition with a similar temporal dynamics and concentration dependence. Neither growth inhibition nor membrane depolarization were caused by the weak acid properties of IAA. The membrane depolarization was only maintained over time at higher concentrations of IAA (100, 1000nM).

RGI requires intracellular accumulation of auxin, but not active transport across the plasma membrane *per se* (Fendrych et al., 2018). On the other hand, previous studies reported the importance of AUX1-mediated influx and PIN2-mediated efflux in the auxin-induced membrane depolarization in root hairs (Dindas et al., 2018; Paponov et al., 2019).

To investigate this contradiction, we increased the intracellular auxin concentration by applying N-1-naphthylphthalamic acid (NPA), an auxin efflux inhibitor that was shown to inhibit root growth rapidly (Fendrych et al., 2018). In our setup NPA only tended to rapidly inhibit root growth (Fig.S2a,b). However, this trend was significant in the steady state response (Fig.S2e). No recordable depolarization was observed neither during the rapid response (Fig.S2c,d) nor during the steady state (Fig.S2f,g), which demonstrated that auxin induced depolarization and RGI can be uncoupled. The reason for the lack of observable depolarization might be that the auxin accumulation caused by NPA is not as sudden as the external IAA treatment. On the other hand, the treatment with NPA significantly impaired the depolarization by both 10nM and 100nM IAA (Fig.2a,b). RGI was only significant at 100nM IAA (Fig.2c,d). Furthermore, during the steady state no depolarization was observed in the presence of NPA (Fig.S2f,g) showing that auxin efflux might be essential for the maintenance of the depolarized state but not essential for the depolarization initiation.

To decipher in more detail the role of auxin transport in membrane depolarization, we genetically perturbed auxin influx, efflux and both influx and efflux using *aux1, pin2* and *aux1pin2* mutants, respectively. Firstly, in rapid response to 10nM IAA, disruption of active IAA influx of IAA in *aux1* resulted in no depolarization (Fig.S2h-j) correlated with the absence of RGI over 5 minutes of treatment (Fig.S2k,l). On the other hand, the *pin2* mutant, impaired in IAA efflux, showed no difference compared to control response in both rapid depolarization (Fig.S2h-j) and RGI (Fig.S2k,l). The simultaneous disruption of IAA influx and efflux in *aux1pin2* resulted in a similar response to *aux1* with no recordable rapid responses (Sup.2h-l). Secondly, to investigate if these results were a consequence of an impaired transport or the lack of the transporters themselves, we saturated tissues with 100nM IAA. In these conditions, *aux1* and *pin2* displayed statistically significant depolarizations, similarly to Col0, while *aux1pin2* only displayed a depolarization which was non significant after 5 minutes of treatment (Fig.2e,f). These responses were correlated with RGI (Fig.2g,h). It is worth highlighting that *aux1* took slightly longer to reach its maximum depolarization. Thirdly, in the steady state experiment with 1000nM IAA, we observed statistically non-significant depolarization tendencies in all three mutants while the control roots were significantly depolarized (Fig.2i). On the other hand, root growth was inhibited in all three mutants and the presence of the *pin2* mutation caused a significant hypersensitivity to IAA (Fig.2j). Together these results showed that AUX1-mediated IAA influx and PIN2-mediated efflux contributed to the maintenance of membrane depolarization, possibly by a constitutive cycling of IAA in and out of cells and the associated proton fluxes. In our experiments, the PIN2 mediated efflux was not required for rapid membrane depolarization, a result contradicting previous studies conducted with 10µM IAA in root hairs showing that both *aux1* and *pin2* mutants are impaired in the membrane potential response (Dindas et al., 2018; Paponov et al., 2019). To put these observations in perspective, these measures were obtained from trichoblasts which might react differently than primary root epidermis. Moreover, in *pin2*, the length and density of root hairs are diminished (Rigas et al., 2013) and thus, could be impaired in auxin response. Finally, at physiological IAA concentrations, AUX1-mediated auxin influx is required for rapid membrane depolarization. Nevertheless, depolarization can be achieved even in the *aux1* mutant with high IAA concentrations when IAA can diffuse into cells. This shows that the depolarization response is not caused by AUX1-mediated IAA influx but by the presence of auxin in the cell and thus, pinpoint towards an intracellular IAA receptor.

The *Arabidopsis thaliana* TIR1/AFB auxin coreceptor, encoded by six paralogous genes TIR1 and AFB1-5 (Dharmasiri et al., 2005), has been separately reported as being involved in both rapid RGI and membrane depolarization by auxin. The *tir1–1/afb2–1/afb3–1* mutant showed a reduced rapid RGI (Fendrych et al., 2018; Scheitz et al., 2013) and decreased depolarization in trichoblast cells (Dindas et al., 2018). Recently, the so-far enigmatic AFB1 paralogue was revealed to play an important role in the rapid root growth response to auxin (Prigge et al., 2020). We confirmed the lack of rapid root inhibition response to auxin in *afb1-3* (Fig.3a,b). At the same time, the *afb1-3* mutant showed a total lack of immediate membrane depolarization response to auxin, even when high IAA concentration was applied (Fig.3c,d,e). Interestingly, after 5 minutes of treatment, *afb1-3* cells initiated a trend for a mild depolarization (Fig.3c). In the steady state, *afb1-3* showed a slightly stronger depolarization in response to IAA (Fig.3f) while the root growth was inhibited albeit less than Col0 (Fig.3g). This might indicate a delay in the mutant’s response to auxin and highlights the crucial role for AFB1 in the earliest auxin response.

Further, we used the orthogonal ccvTIR1 - cvxIAA receptor - auxin pair (Uchida et al., 2018) in which all the native TIR1/AFB paralogues are shunted and only the synthetic ccvTIR1 receptor is active when synthetic cvxIAA is applied. The cvxIAA was shown to rapidly inhibit growth of ccvTIR1 roots. However, a several minute delay in the growth response was recorded when compared to the IAA - wild type root combination (Fendrych et al., 2018). We used the pico-cvxIAA synthetic auxin that is active in lower doses compared to cvxIAA (Yamada et al., 2018). In ccvTIR1 roots, pico-cvxIAA did not trigger an observable rapid depolarization (Fig.3h,i). In the observation window (0 to 10 minutes), pico-cvxIAA treated roots initiated a trend to inhibit root growth around 7 minutes of treatment (Fig.3j,k). This trend was significant after 15 minutes. (Fig.S3a). In the steady state, the only observable effect of pico-cvxIAA on membrane potential was a hyperpolarization of Col0’s root membranes when treated with 1000nM (Fig.S3b). These results revealed that in steady state, RGI and depolarization can be uncoupled and tend to point toward non-rapid signaling mechanisms. Col0 was not affected by both 100 and 1000nM of pico-cvxIAA while ccvTIR1 showed a strong RGI (Fig.S3c).

Finally, as the ccvTIR1 receptor is expressed in the *tir1/afb2* mutant background, we analyzed the response of this line to IAA to address to what extent the TIR1 and AFB2 are necessary for the rapid membrane depolarization. We observed an instantaneous but weaker and slowly increasing IAA-induced rapid depolarization (Fig.3h,i) and impaired RGI (Fig.3j,k) compared to control. In the steady state, no significant depolarization was observed in response to high IAA concentration (Fig.S3b). However, with this treatment, the mutant displayed a similar RGI as Col0 (Fig.S3c).

Altogether, these results clarified the role of the TIR1/AFB family of auxin receptors in auxin-triggered membrane depolarization: The initial instantaneous depolarization and RGI strictly depend on AFB1. When AFB1 is missing, as in the *afb1-3* mutant or in the pico-cvxIAA - ccvTIR1 combination, the depolarization response is delayed or absent. The ccvTIR1 receptor itself is unable to trigger the full rapid auxin response that involves membrane depolarization, which explains the delay in root growth inhibition observed before (Fendrych et al., 2018), highlights the importance of AFB1 and also demonstrates the role of membrane depolarization in root growth inhibition. At the same time, the other paralogues such as TIR1 and AFB2 clearly participate in the rapid response, as shown by the decreased intensity of depolarization in the mutant *tir1/afb2* background, as well as by the data in root hairs (Dindas et al., 2018; Paponov et al., 2019). The steady state depolarization was not always fully correlated with the mutant’s growth response, similarly to perturbations in auxin transport, indicating that the maintenance of the depolarization is a very sensitive readout of genetic perturbation. The inability of the cvxIAA-ccvTIR1 pair to trigger long-term membrane depolarization might be caused by the lack of cvxIAA transport by the auxin carriers.

We showed that the AFB1 receptor is essential for the rapid initiation of IAA-induced depolarization and RGI. Nevertheless, the relevance of the auxin-induced membrane depolarization for actual physiological processes remains unclear. During gravitropic bending, the lower root side accumulates IAA, leading to differential root elongation between lower and upper side resulting in root bending. Using impaling probes, it was shown that the lower epidermis of gravitropically bending maize roots was depolarized while the upper side showed hyperpolarization in comparison to the situation during vertical growth (Behrens et al., 1985; Ishikawa and Evans 1990). To test the significance of auxin induced depolarization and growth inhibition in the gravitropic root response, we established imaging of membrane potential during the gravitropic response.

The staining with DISBAC_2_(3) did not interfere with the gravitropic response when grown on the agar surface (Fig.S4a). However, when the stained roots were grown between the agar and the imaging coverglass, the staining slowed down the gravitropic response (Fig.S4b). To avoid any bias in our measurements, we restricted our observation of membrane potential response to the first 10 minutes of the gravitropic response.

Before gravistimulation, the membrane potential profiles of left and right sides of the root were similar (Fig.4a). Two minutes after 90° rotation of the seedlings, we observed a clear trend of the lower side to be more depolarized in the elongation zone (150-300µm from the quiescent center, Fig.4b) while the upper side tended to be hyperpolarized. This non-significant tendency was also observed by measuring the fluorescence of the transition zone (Fig.4c). However, after 10 minutes this tendency inverted. During gravitropic bending, the concentration of IAA in the lower side increases approximately two-fold (Band et al., 2012). This concentration rise is probably at the detection limit of DISBAC_2_(3) staining and thus produces tendencies and not statistically significant results. Moreover, this tendency is in accordance with the literature (Behrens 1985; Ishikawa and Evans 1990).

To test of whether modulation of membrane polarization can prevent auxin-triggered depolarization during gravitropism and in turn interfere with gravitropic bending, we hyperpolarized the root membranes by exogenous application of FC (Fig.S1d,e). This treatment, however, did not prevent auxin-induced depolarization observed at the lower side after 2 minutes of gravistimulation (Fig.4c), producing the exact same tendency as the mock treatment. In agreement with this, the root bending response was similar to control in the initial stages but led to overbending in the late phase of gravitropic response (Fig.S4c). This demonstrates that auxin is still able to trigger a depolarization and growth response even in a root hyperpolarized by FC.

We demonstrated that *afb1-3* plants showed neither rapid depolarization nor rapid RGI in response to auxin. We therefore tested the gravitropism-induced depolarization of the *afb1-3* mutant. No difference or tendency was observed between polarization profiles obtained before (Fig.4a,c) and after 2 minutes of gravistimulation (Fig.4b,c). Nevertheless, 10 minutes after gravistimulation, the *afb1-3* showed a tendency towards depolarization in the lower side of the roots (Fig.3c-e), which was not observed in the wild type plants.

It was previously shown that despite the defects in auxin response, the *afb1-3* single mutant has a normal gravitropic response (Prigge et al., 2020). In a low-resolution gravitropic experiment, we obtained similar results where the *afb1-3* gravitropism was not significantly different from the behavior of the control (Fig.4d). However, *afb1-3* being mostly impaired in the earliest stages of auxin response, we optimized high spatiotemporal resolution imaging of early root gravitropism (Fig.S4d) and quantified the root bending angle using an unbiased semi-automated image analysis workflow. Using this approach, the control plants initiated gravitropic bending within a few minutes after the gravitropic stimulus while the *afb1-3* roots showed an approximately 10-minute delay (Fig.4e,f, supplemental movie 2 and 3). These results spectacularly match the defects in membrane depolarization observed during external application of IAA as well as the growth inhibition impairment observed in the *afb1-3* mutant. Intriguingly, the *afb1-3* mutant phenotype resembles the behavior of the roots lacking the calcium channel CNGC14 (Shih et al., 2015). On the other hand, the rapidity of the gravitropic response in the control roots confirms that rapid responses to auxin are indeed happening and are relevant for plant cell responses to the internal auxin fluxes.

## Conclusion

We established a toolbox to dynamically visualize membrane potential *in vivo* in *Arabidopsis thaliana* roots by combining the DISBAC_2_(3) fluorescent probe with microfluidics and vertical stage microscopy. This way we show that auxin-induced membrane depolarization tightly correlates with rapid RGI and that the cells of the transition zone/early elongation zone are the most responsive to auxin. Further, we demonstrate that auxin cycling in and out of the cells through AUX1 influx and PIN2 efflux is *per se* not essential for membrane depolarization and rapid RGI but is a facilitator of these responses. The rapid membrane depolarization by auxin instead strictly depends on the AFB1 auxin receptor, while the other TIR1/AFB paralogues contribute to this response. The lack of membrane depolarization in the *afb1* mutant explains the lack of the immediate root growth inhibition. Finally, we show that AFB1 is required for the rapid depolarization and rapid growth inhibition of cells at the lower side of the gravistimulated root. Taken together, these results mark a major step in understanding the physiological significance of membrane depolarization for the gravitropic response of the root and clarify the role of AFB1 as the receptor central for rapid auxin responses, adding another piece to the puzzle in understanding the biology of the phytohormone auxin.

## Acknowledgements

This work was supported by the European Research Council (Grant No. 803048), Charles University Primus (Grant No. PRIMUS/19/SCI/09), and in the initial stages by the Czech Science Foundation (GAČR Grant No. 18-10116Y). Sergey Shabala acknowledged support from the Australian Research Council and China National Distinguished Expert Project (WQ20174400441).

The authors thank Mark Estelle and Mike Prigge for Arabidopsis seeds, Jiří Friml for the initial *aux1pin2* cross, Jack Merrin for guidance in microfluidics.

## Author contributions

NBCS and MF conceived the project, performed the experimental work, analyzed, and interpreted the data. PY and SS conceived the impaling electrode membrane potential experiment and PY measured and analyzed the data. DK, ZS and MF conceived the microfluidics chip design and DK fabricated and optimized it. NBCS, MF and SS wrote the manuscript.

## Material and methods

### Plant material and growth conditions

Wild-type *Arabidopsis thaliana* ecotype Columbia (Col0) and the following transgenic lines were used in this study: *aux1* (SALK_020355), *pin2* (NASC_N16706), *aux1pin2* (SALK_020355/SALK_091142), ccvTIR1 (*tir1-afb2*, Uchida et al., 2018) and *afb1-3* (SALK_ SALK_070172, Salvadi-Goldstein et al., 2008).

Genotypes were verified by PCR-genotyping using the following primers. SALK LB1.3 primer (ATTTTGCCGATTTCGGAAC) and *aux1*-R (AGCTGCGCATCTAACCAAGT) (and the aux1-L primer), *pin2*-R (AAGCACCAAAGACTATAACTA) and the PIN2 L primer, *afb1-3*-RP (GCAACAGCTTCAAGACCTTTG) and afb1-3-LP (AACGGAAGACTAGGAAGCGAG). ccvTIR1-R was verified by its ability to react to pico-cvxIAA. *pin2* single mutant has a single nucleotide (G to A) insertion which was verified by comparison of Col0 and *pin2* sequencing of the PCR product obtained with the primers: PIN2-R (AAGCACCAAAGACTATAACTA) and PIN2-F(CAACGCGAAGAATGCTATGA).

Seeds were surface sterilized by chlorine gas for 2 hours (Lindsey et al., 2017). Seeds were sown on 1% (w/v) agar (Duchefa) with ½ Murashige and Skoog (MS, Duchefa, 1 % (w/v) sucrose, adjusted to pH 5.8 with KOH 1M, and stratified for 2 days at 4°C. Seedlings were grown vertically in a growth chamber with 23°C by day (16h), 18°C by night (8h), 60% humidity, light intensity of 120 µmol photons m^-2^ s^-1^.

### Pharmacological treatments

Treatments were prepared using the following chemicals: 3-indoleacetic acid (IAA, 10mM stock in 96% ethanol, Sigma Aldrich), N-1-naphthylphthalamic acid (NPA, 10mM stock in DMSO, Sigma Aldrich), Carbonyl cyanide m-chlorophenyl hydrazone (CCCP, 10mM stock in DMSO, Sigma Aldrich), Fusicoccin (FC, 1mM stock in 96% ethanol, Sigma Aldrich), 5-Adamantyl-IAA (pico-cvxIAA, 10mM stock in DMSO, TCI Chemicals).

### Microfluidic chip description and manufacturing

Microfluidics experiments were conducted using a closable single-layer PDMS silicone chip (Fig.S1g). The chip contained two inlet channels with dimensions of 200 × 50 µm (w×h) and two channels accommodating the growing roots of 1000 × 100 × 20000 µm (w×h×l). To facilitate the sealing of the PDMS and the coverslip, the PDMS resin (Sylgard 184, Dow Corning, USA) was prepared by mixing 15:1 parts of PDMS base: curing agent, resulting in a stickier surface. The master for PDMS casting was made from a two-component polyurethane resin (PUR, F32, Axson, Czech Republic) that consists of Polyol (main base) and Isocyanate (curing agent). These were mixed in a 1:1 volumetric ratio and cured for 15 minutes. To prepare the master, 4 ml of resin was poured on a plexiglass plate with microfluidic structures identical to those in the final chip. These structures were sculptured by micromachining using a micro-milling machine (GV21, GRAVOS, Czechia).

### Microfluidic experiments

Five-day-old seedlings were transferred to the chip (Fig.S1g) and enclosed by careful placement of a pre-cleaned microscopic glass coverslip (117µm thick) on the PDMS cast. The closed chip was then placed in a home-made plexiglass holder with screws ensuring tight sealing between the PDMS layer and the coverslip (Fig.S1h). This setup was then placed on the vertical microscope stage for 20 minutes to recover before imaging. A constant flow of 3 µL/minute was maintained using a piezo electric pressure controller (OBI1, Elveflow, France) coupled with micro-flow sensors (MFS2, Elveflow, France) and the dedicated Elveflow software to control both recording and the flow/pressure feedback. To switch between control and treatment solutions, we set the desired solution flow to 3 +/- 0.01µL/minute and the other to 0 +/- 0.01µL/min.

### Membrane potential recording

Conventional 1M KCl-filled Ag/AgCl microelectrodes with a tip diameter of 0.5μm were used (Shabala et al., 2005). The electrodes were connected to the MIFE amplifier (University of Tasmania, Hobart, Australia) and voltage reading recorded with the CHART software (see Shabala et al., 2006 for details). Seedlings were immobilized in 0.5 g/L MES solution containing various amounts of KCl (ranging from 10μM to 100mM). After 30 min of exposure, electrodes were impaled into epidermal cells in the mature root zone, and membrane potential values recorded for at least 20 sec. At least 5 individual plants were measured for each treatment, with 4 to 6 cells impaled in each (n = [20-30]).

For membrane potential measurement using DISBAC_2_(3) fluorescence, steady state experiments were conducted using 5-day old seedlings transferred to ½ MS, 1% (w/v) sucrose in 1% agar (Duchefa) and 15µM DISBAC_2_(3) (Sigma-Aldrich). Seedlings and staining/treatment media were then placed into a custom 3D printed chambered coverglass (24 x 60mm) and treated for 20 minutes before imaging every 10 minutes for 40 minutes. Control media contained the same amount of the chemical solvent added to the treatment mediums (½ MS, DMSO or ethanol 96%). For microfluidics experiments, both control and treatment solutions were prepared by adding 1% (w/v) sucrose and then 20µM DISBAC_2_(3) to a pre-solution, ensuring that both mediums contained the same amount of sucrose and dye. This pre-solution was then split in two and chemical(s) were added to the treatment solution.

The same volume of chemical solvent was added to the control solution (½ MS, DMSO or ethanol 96%). Two seedlings were transferred to the two channels pre-filled with control solution. After closing the chip, the channels were constantly flushed with control solution and placed on the vertical microscopy stage for 20 minutes before imaging. Both seedlings were then imaged every 30 seconds. We recorded 6 minutes of control and 12 minutes of treatment (except specified otherwise).

### Analysis of the gravitropic response

The low-resolution analysis of gravitropic response was performed using a vertically placed flatbed scanner (Perfection V600, Epson). 5-day old seedlings were transferred to plates containing the desired media and let to recover in a growth chamber. After an hour, plates were turned 90° and imaged every 30 minutes.

For high-spatiotemporal analysis of gravitropism, 4-day old seedlings were placed onto a thin layer of ½ MS medium placed inside a custom 3D printed chambered coverglass (24 x 50mm). The seedlings were let to recover vertically for at least 30 minutes before gravistimulation. In this setup, the roots were growing unobstructed on the surface of the agar and the imaging was performed through the coverglass and the agar (Fig.S4d). Three roots of control and mutant plants were imaged at the same time every 1 minute for 40 minutes.

### Microscopic imaging

Imaging was performed using a vertical stage (von Wangenheim et al., 2017) Zeiss Axio Observer 7 coupled to a Yokogawa CSU-W1-T2 spinning disk unit with 50 µm pinholes and equipped with a VS-HOM1000 excitation light homogenizer (Visitron Systems). Images were acquired using the VisiView software (Visitron Systems) and Zen Blue (Zeiss) for fig. 4e, f. We used the Zeiss Plan-Apochromat 20x/0.8 and Plan-Apochromat 10x/0.45 objectives. DISBAC_2_(3) was excited with a 515nm laser and the emission was filtered by a 535/30nm band pass filter. Signal was detected using a PRIME-95B Back-Illuminated sCMOS Camera (1200 x 1200 px; Photometrics) or Orca Flash 4.0 V3 (2048 x 2048 px; Hamamatsu) for fig. 4e, f.

**Fig 1:**
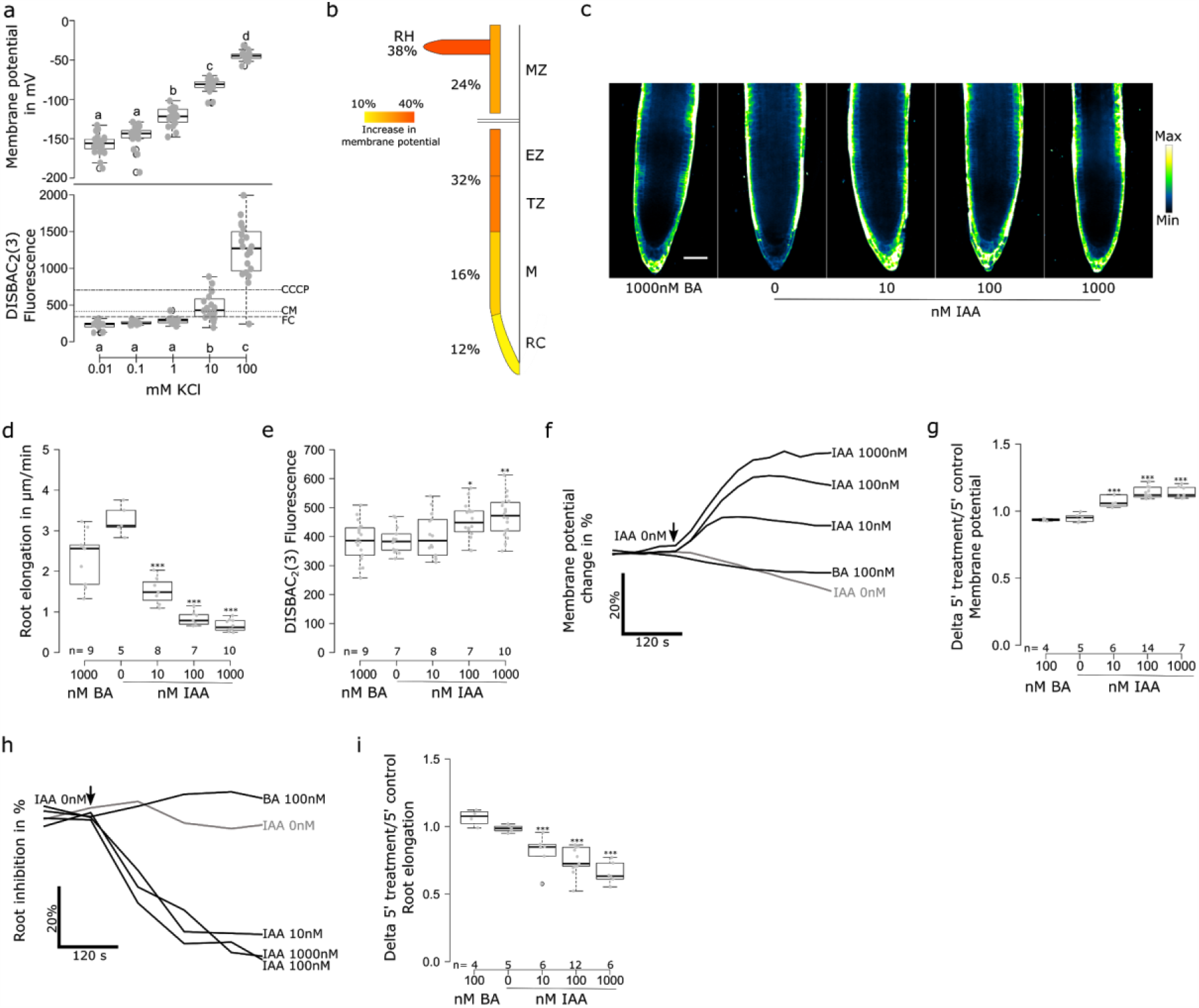
Auxin induces depolarization and root growth inhibition in *Arabidopsis thaliana* root tip. a) Comparison of impaling probe membrane potential measurements (in mV) and DISBAC_2_(3) fluorescence (arbitrary unit) in response to a KCl gradient. Dashed lines represent DISBAC_2_(3) median fluorescence values of control (½ MS), CCCP and FC treatments, indicating that DISBAC_2_(3) detection range covers the physiological range of membrane potential. b) Root epidermis membrane potential response to 100nM IAA in the root cap (RC), meristematic zone (M), transition/elongations zone (TZ and EZ) and mature zone (MZ). Percentages represent the delta of DISBAC_2_(3) fluorescence between control media and 5 minutes of treatment. n= 5 individual roots. c-i) Effects of an IAA concentration gradient on c) steady state root tip membrane potential and d) primary root elongation (µm/minute). f) rapid membrane potential response mean change (in %) over 5 minutes of treatment and g) ratio of 5 minutes treatment over 5 minutes control media. h) rapid root elongation response, average change (in %) over 5 minutes of treatment and i) delta of 5 minutes treatment over 5 minutes control media. Steady state corresponds to the fluorescence of roots after 20 minutes of treatment in agar medium and root elongation measured for 20 minutes. Rapid response corresponds to roots treated in microfluidics and measured every 30 seconds. Application of treatments is indicated by a black arrow. For f) and h), standard errors were not added to simplify reading. c) representative microscopy images look up table indicated on the right, scale bar = 50µm. n= is indicated on figures. ***: p-value<0.0005, *: p-value<0.05.

**Figure 2:**
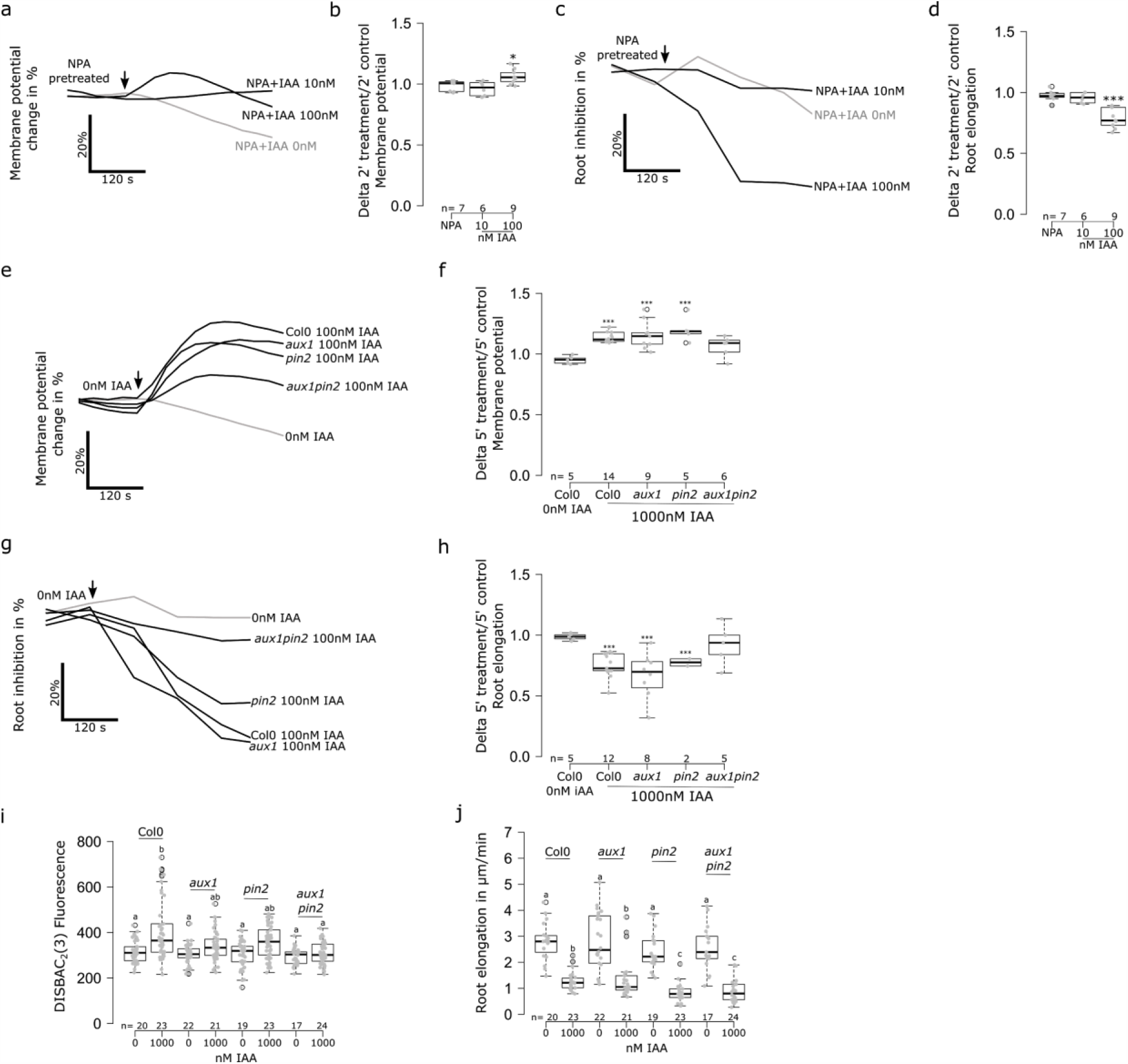
Auxin transport through PIN2 and AUX1 is not essential for rapid membrane depolarization. a-d) Effect of NPA on the rapid response to 0, 10 or 100nM IAA; roots pretreated with 20µM NPA. a) Membrane potential average change (in %) over 5 minutes of treatment and b) delta of 2 minutes treatment over 2 minutes control media. c),d) Root elongation over 5 minutes of treatment. c) average change (in %) over 5 minutes of treatment and d) delta of 2 minutes treatment over 2 minutes control media. e-g) Effect of 10 nM IAA on Col0, *aux1, pin2* and *aux1pin2* the rapid membrane potential response. e) Membrane potential fluorescence in the transition zone cells. f) average change (in %) over 5 minutes of treatment and g) delta of 5 minutes treatment over 5 minutes control media. h) Root elongation rapid response over 10 minutes of treatment; average change (in %) over 5 minutes of treatment and i) delta of 5 minutes treatment over 5 minutes control media. j) Effect of 1000 nM IAA on Col0, *aux1, pin2* and *aux1pin2* on the steady state response of root tip membrane potential and k) primary root elongation (µm/minute). Steady state corresponds to the fluorescence of roots after 20 minutes of treatment in agar medium and root elongation measured over 20 minutes. Rapid response corresponds to roots treated in microfluidics and measured every 30 seconds. Application of treatments is indicated by a black arrow. For a, c, f and h, standard errors were not added to simplify reading. In e) the look up table is indicated on the right. n= is indicated on figures. ***: p-value<0.0005.

**Figure 3:**
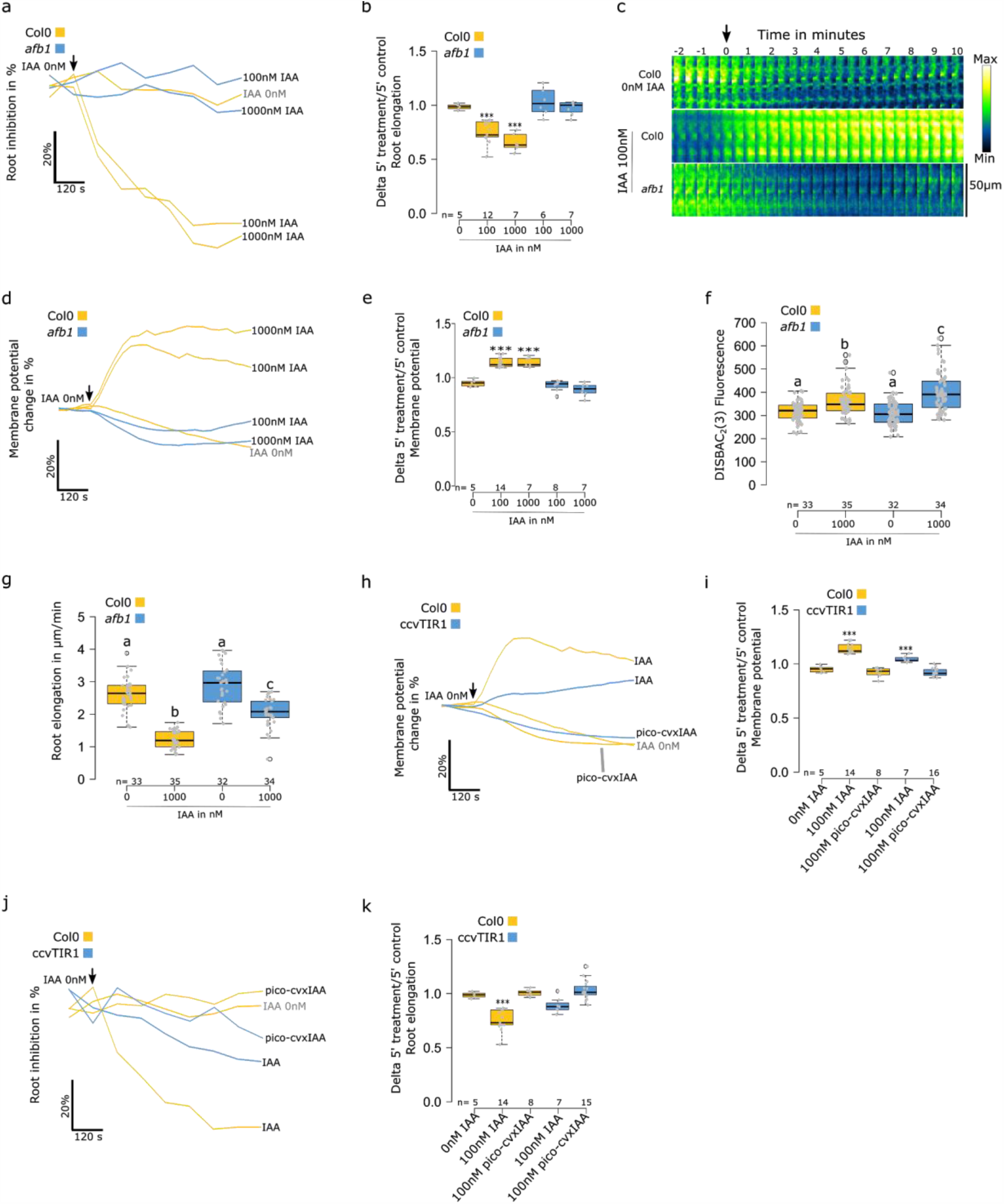
AFB1 triggers rapid membrane depolarization in response to IAA. a-g) Effect of IAA on Col0 and *afb1-3*. a) Rapid root elongation response, average change (in %) over 10 minutes of treatment and b) delta of 5 minutes treatment over 5 minutes control media. c-e) membrane potential rapid response change over 10 minutes of treatment. c) Representative images of membrane potential reporter in time. d) average change (in %) over 10 minutes of treatment and i) delta of 5 minutes treatment over 5 minutes control media. f) Steady state response of root tip membrane potential and g) primary root elongation (µm/minute). h-k) Effect of IAA and pico-cvxIAA on Col0 and ccvTIR1. h) Rapid membrane potential response; average change (in %) over 10 minutes of treatment and i) delta of 5 minutes treatment over 5 minutes control media. j) Root elongation rapid response over 10 minutes of treatment; average change (in %) over 5 minutes of treatment and k) delta of 5 minutes treatment over 5 minutes control media. Steady state corresponds to the fluorescence of roots after 20 minutes of treatment in agar medium and root elongation measured over 20 minutes. Rapid response corresponds to roots treated in microfluidics and measured every 30 seconds. Application of treatments is indicated by a black arrow. For a, c, f and h, standard errors were not added to simplify reading. c) Fluorescence intensity look up table is indicated on the right. n= is indicated on figures. ***: p-value<0.0005, *: p-value<0.05.

**Figure 4:**
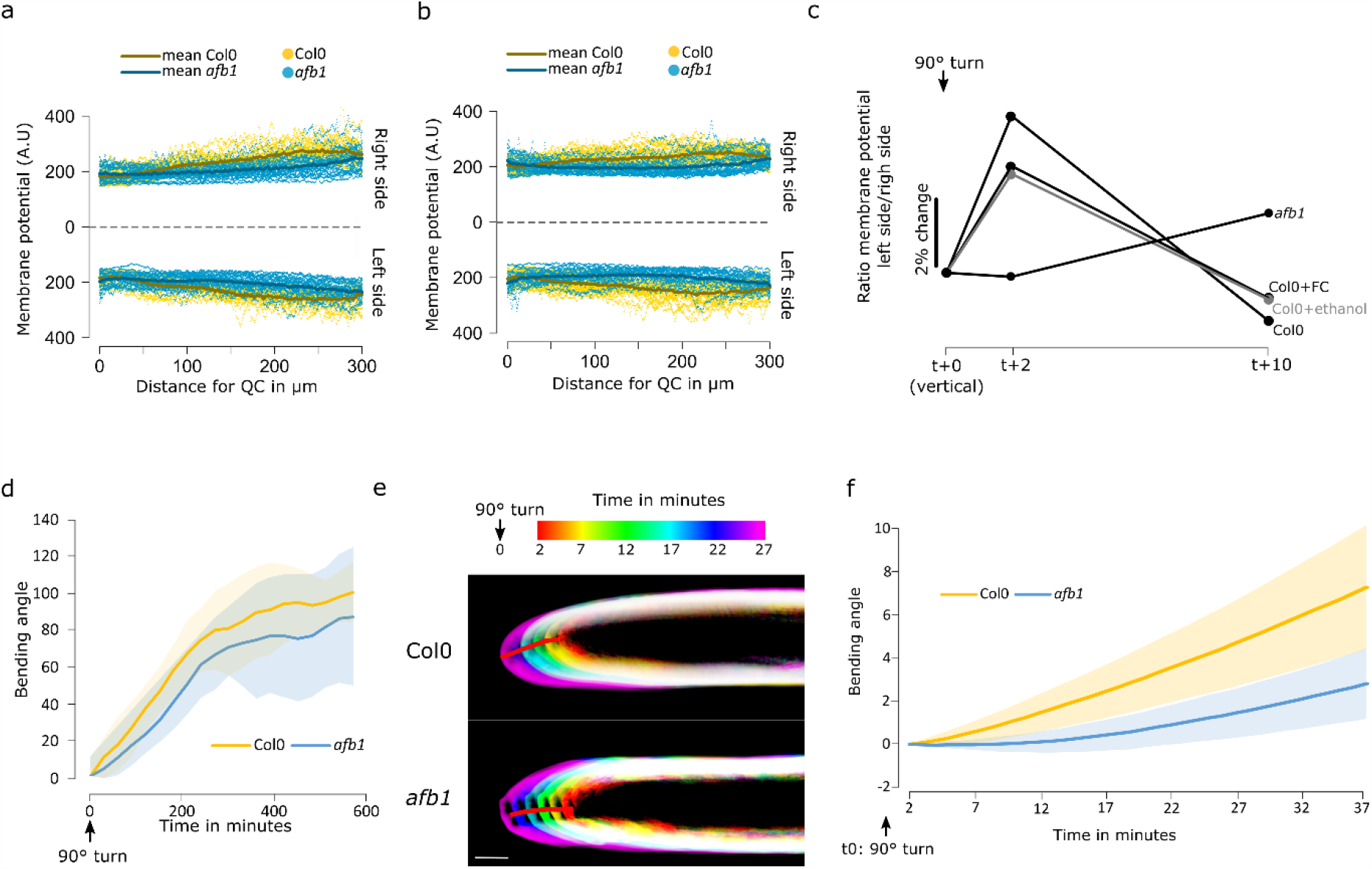
AFB1-induced depolarization drives the early stages of root gravitropic response. a),b) Membrane potential of individual roots and average fluorescence profiles from quiescent center (0µm) to elongation zone (300µm) of Col0 and *afb1-3* during a) vertical growth and b) after 2 minutes of 90° gravistimulation. n= [22-25] individual roots. c) Ratios of membrane potential from lower/upper side of the root transition zone. Ratios were measured at t+0 (vertical growth), t+2 (2 minutes of 90° gravistimulation) and t+10 (10 minutes of 90° gravistimulation) from root in control media (Col0), ethanol (Col0 + ethanol) and 5µM fusicoccin (Col0 + FC). Ratios are expressed in % of changes from t+0. n= [22-25] individual roots. d) Bending angle during 550 minutes after 90° gravistimulation of Col0 and *afb1-3*. Images were taken every 30 minutes. n= [22-25] individual roots. e),f) High spatio-temporal resolution imaging of Col0 and *afb1-3* response to gravistimulation e) Temporal color code images of representative Col0 and *afb1-3* roots. f) Quantification of the root tip angles. n= [8-12] individual roots.

### Image analysis

All microscopy image analysis were conducted using the software ImageJ Fiji (Schindelin et al., 2012). DISBAC_2_(3) fluorescence was measured using the segmented line selection tool (20 pixels width). Transition zone was defined as starting at the first epidermis cell (from the quiescent center) that doubled in length compared to the previous one and ending 6 cells shootward (approximately 100µm). The selection line was placed at the intersection of epidermis and cortex cells. Fluorescence was measured in both sides of the root.

Root growth rate was measured with the FIJI plugin Correct 3D drift (Parslow et al., 2014) by stabilizing the drift of the root tip. From the drift exported file, distance between root tip positions in consecutive frames was calculated using the formula to calculate distance between 2 sets of coordinates in space.

Root bending angle in the low-resolution scanner experiments was measured after transforming the roots into strings of pixels. The angle between the root tip pixel and the 10^th^ pixel before the root tip was then calculated for every timeframe.

Root bending during high-resolution microscopy experiments was measured by calculating the intersection angle between a line cutting the root tip in two and a perpendicular line to the line cutting the shoot side root in two.

To produce homogeneous measurements and unbiased analysis, measurements of root growth and root bending in scanner and microscope experiments were semi-automatized to limit human-picture interaction/interpretation using ImageJ macros and R scripts (R Core Team, R Foundation for Statistical Computing, Austria, R Cran v3.5.3 and R Studio v1.1.463). The following R packages were used: LearnGeom (Briz-Redón and Serrano-Aroca 2018), spdep (Bivand and Wong 2018), stringr (Wickam et al., 2019, https://CRAN.R-project.org/package=stringr), dplyr (Wickam et al., 2020, https://CRAN.R-project.org/package=dplyr), REdaS (Hatzinger et al 2014), reshape2 (Wickam et al., 2007), matrixstats (Bengtsson et al., 2020; https://CRAN.R-project.org/package=matrixStats).

### Graphics and statistical analysis

Graphics and statistical analysis were performed using the R software. As our samples were under 40 individuals and then, unlikely to follow a normal distribution and respect the equality of variances, non-parametric comparison tests were used with 95% confidence using the nparcomp R package (Konietshke et al 2015). To compare two sets of samples we used a non-parametric Student test (npar.t.test function). To compare several samples, we used non-parametric multiple contrast tests (mctp function). We adjusted the type of comparison by using either a Dunnett (every sample compared to a control) or Tukey (every sample between them) contrast method according to the statistical analysis desired.

Every steady state experiment was conducted two to four times using different seedlings sown on different days. All the conditions presented in one boxplot were imaged and measured at the same time. For fluorescence, raw arbitrary units are presented.

For microfluidics experiments, the results correspond to a minimum of 4 individual chips containing two seedlings. The measurements were conducted using seedlings sown on at least 2 different days. As microfluidics experiments are independent to each other (control and treatment for one genotype in one experiment) we took the liberty to reuse, on different graphics and analysis, the data collected with Col0 and the following treatment: CM to CM, CM to IAA 10nM, CM to IAA 100nM. Both fluorescence and root elongation data presented have been normalized to reflect changes in percentages. The normalization for one individual root was performed by dividing each value for each time point by the mean of 5 minutes of control measurements (every 30 seconds, otherwise specified). This ratio was then multiplied by 100 to obtain percentages. The ratio shown in boxplot figures represents the mean of 5 minutes of treatment measurements (every 30 seconds, otherwise specified) by the mean of 5 minutes (except specified otherwise) of control measurements.

## References

Band, L.R., Wells, D.M., Larrieu, A., Sun, J., Middleton, A.M., French, A.P., Brunoud, G., Sato, E.M., Wilson, M.H., Peret, B., Oliva, M., Swarup, R., Sairanen, I., Parry, G., Ljung, K., Beeckman, T., Garibaldi, J.M., Estelle, M., Owen, M.R., Vissenberg, K., Hodgman, T.C., Pridmore, T.P., King, J.R., Vernoux, T., Bennett, M.J., 2012. Root gravitropism is regulated by a transient lateral auxin gradient controlled by a tipping-point mechanism. Proceedings of the National Academy of Sciences 109, 4668–4673.https://doi.org/10.1073/pnas.1201498109

Bates, G.W., Goldsmith, M.H., 1983. Rapid response of the plasma-membrane potential in oat coleoptiles to auxin and other weak acids. Planta 159, 231–237. https://doi.org/10.1007/BF00397530

Baunsgaard, L., Fuglsang, A.T., Jahn, T., Korthout, H.A., de Boer, A.H., Palmgren, M.G., 1998. The 14-3-3 proteins associate with the plant plasma membrane H+-ATPase to generate a fusicoccin binding complex and a fusicoccin responsive system. Plant J 13, 661–671. https://doi.org/10.1046/j.1365-313x.1998.00083.x

Behrens, H.M., Gradmann, D., Sievers, A., 1985. Membrane-potential responses following gravistimulation in roots of Lepidium sativum L. Planta 163, 463–472. https://doi.org/10.1007/BF00392703

Bivand, R.S., Wong, D.W.S., 2018. Comparing implementations of global and local indicators of spatial association. TEST 27, 716–748. https://doi.org/10.1007/s11749-018-0599-x

Briz-Redón, Á., Serrano-Aroca, Á., 2018. Novel pedagogical tool for simultaneous learning of plane geometry and R programming. RIO 4, e25485. https://doi.org/10.3897/rio.4.e25485

Dharmasiri, N., Dharmasiri, S., Weijers, D., Lechner, E., Yamada, M., Hobbie, L., Ehrismann, J.S., Jürgens, G., Estelle, M., 2005. Plant Development Is Regulated by a Family of Auxin Receptor F Box Proteins. Developmental Cell 9, 109–119. https://doi.org/10.1016/j.devcel.2005.05.014

Dindas, J., Becker, D., Roelfsema, M.R.G., Scherzer, S., Bennett, M., Hedrich, R., 2020. Pitfalls in auxin pharmacology. New Phytol 227, 286–292. https://doi.org/10.1111/nph.16491

Dindas, J., Scherzer, S., Roelfsema, M.R.G., von Meyer, K., Müller, H.M., Al-Rasheid, K.A.S., Palme, K., Dietrich, P., Becker, D., Bennett, M.J., Hedrich, R., 2018. AUX1-mediated root hair auxin influx governs SCFTIR1/AFB-type Ca2+ signaling. Nature Communications 9. https://doi.org/10.1038/s41467-018-03582-5

Evans, M.L., Ishikawa, H., 1994. Responses of Arabidopsis roots to auxin studied with high temporal resolution: Comparison of wild type and auxin-response mutants. Planta 194, 215–222. https://doi.org/10.1007/BF01101680

Evans, M.L., Mulkey, T.J., Vesper, M.J., 1980. Auxin action on proton influx in corn roots and its correlation with growth. Planta 148(5), 510–512. https://doi.org/10.1007/BF00552667

Felle, H., Peters, W., Palme, K., 1991. The electrical response of maize to auxins. Biochimica et Biophysica Acta (BBA) - Biomembranes 1064, 199–204. https://doi.org/10.1016/0005-2736(91)90302-O

Fendrych, M., Akhmanova, M., Merrin, J., Glanc, M., Hagihara, S., Takahashi, K., Uchida, N., Torii, K.U., Friml, J., 2018. Rapid and reversible root growth inhibition by TIR1 auxin signalling. Nature Plants 4, 453–459. https://doi.org/10.1038/s41477-018-0190-1

Goering, H., Stahlberg, R., 1979. Depolarization of transmembrane potential of corn and wheat coleoptiles under reduced water potential and after IAA application. Plant and Cell Physiology 20(3), 649–656. https://doi.org/10.1093/oxfordjournals.pcp.a075851

Hatzinger, R., Hornik, K., Nagel, H., Maier, M.J., 2014. R: Einführung durch angewandte Statistik, 2., aktualisierte Auflage. ed. Pearson, Hallbergmoos/Germany.

Konietschke, F., Placzek, M., Schaarschmidt, F., Hothorn, L.A., 2015. nparcomp: An R Software Package for Nonparametric Multiple Comparisons and Simultaneous Confidence Intervals. Journal of Statistical Software 64. https://doi.org/10.18637/jss.v064.i09

Labady, A., Thomas, D., Shvetsova, T., Volkov, A.G., 2002. Plant bioelectrochemistry: effects of CCCP on electrical signaling in soybean. Bioelectrochemistry 57, 47–53.https://doi.org/10.1016/S1567-5394(01)00175-X

Lindsey, B.E., Rivero, L., Calhoun, C.S., Grotewold, E., Brkljacic, J., 2017. Standardized Method for High-throughput Sterilization of Arabidopsis Seeds. JoVE 56587. https://doi.org/10.3791/56587

Monshausen, G.B., Miller, N.D., Murphy, A.S., Gilroy, S., 2011. Dynamics of auxin-dependent Ca2+ and pH signaling in root growth revealed by integrating high-resolution imaging with automated computer vision-based analysis: Calcium and auxin signaling. The Plant Journal 65, 309–318. https://doi.org/10.1111/j.1365-313X.2010.04423.x

Paponov, I.A., Dindas, J., Król, E., Friz, T., Budnyk, V., Teale, W., Paponov, M., Hedrich, R., Palme, K., 2019. Auxin-Induced Plasma Membrane Depolarization Is Regulated by Auxin Transport and Not by AUXIN BINDING PROTEIN1. Frontiers in Plant Science 9. https://doi.org/10.3389/fpls.2018.01953

Parslow, A., Cardona, A., Bryson-Richardson, R.J., 2014. Sample Drift Correction Following 4D Confocal Time-lapse Imaging. JoVE 51086. https://doi.org/10.3791/51086

Prigge, M.J., Platre, M., Kadakia, N., Zhang, Y., Greenham, K., Szutu, W., Pandey, B.K., Bhosale, R.A., Bennett, M.J., Busch, W., Estelle, M., 2020. Genetic analysis of the Arabidopsis TIR1/AFB auxin receptors reveals both overlapping and specialized functions. eLife 9, e54740. https://doi.org/10.7554/eLife.54740

Renier, M., Tamanini, A., Nicolis, E., Rolfini, R., Imler, J.-L., Pavirani, A., Cabrini, G., 1995. Use of a Membrane Potential-Sensitive Probe to Assess Biological Expression of the Cystic Fibrosis Transmembrane Conductance Regulator. Human Gene Therapy 6, 1275–1283. https://doi.org/10.1089/hum.1995.6.10-1275

Rigas, S., Ditengou, F.A., Ljung, K., Daras, G., Tietz, O., Palme, K., Hatzopoulos, P., 2013. Root gravitropism and root hair development constitute coupled developmental responses regulated by auxin homeostasis in the Arabidopsis root apex. New Phytol 197, 1130–1141. https://doi.org/10.1111/nph.12092

Savaldi-Goldstein, S., Baiga, T.J., Pojer, F., Dabi, T., Butterfield, C., Parry, G., Santner, A., Dharmasiri, N., Tao, Y., Estelle, M., Noel, J.P., Chory, J., 2008. New auxin analogs with growth-promoting effects in intact plants reveal a chemical strategy to improve hormone delivery. Proceedings of the National Academy of Sciences 105, 15190–15195. https://doi.org/10.1073/pnas.0806324105

Schindelin, J., Arganda-Carreras, I., Frise, E., Kaynig, V., Longair, M., Pietzsch, T., Preibisch, S., Rueden, C., Saalfeld, S., Schmid, B., Tinevez, J.-Y., White, D.J., Hartenstein, V., Eliceiri, K., Tomancak, P., Cardona, A., 2012. Fiji: an open-source platform for biological-image analysis. Nat Methods 9, 676–682. https://doi.org/10.1038/nmeth.2019

Senn, A.P., Goldsmith, M.H.M., 1988. Regulation of Electrogenic Proton Pumping by Auxin and Fusicoccin as Related to the Growth of Avena Coleoptiles. PLANT PHYSIOLOGY 88, 131–138. https://doi.org/10.1104/pp.88.1.131

Shabala, S., Demidchik, V., Shabala, L., Cuin, T.A., Smith, S.J., Miller, A.J., Davies, J.M., Newman, I.A., 2006. Extracellular Ca 2+ Ameliorates NaCl-Induced K + Loss from Arabidopsis Root and Leaf Cells by Controlling Plasma Membrane K +-Permeable Channels. Plant Physiol. 141, 1653–1665. https://doi.org/10.1104/pp.106.082388

Shih, H.-W., DePew, C.L., Miller, N.D., Monshausen, G.B., 2015. The Cyclic Nucleotide-Gated Channel CNGC14 Regulates Root Gravitropism in Arabidopsis thaliana. Curr Biol 25, 3119–3125. https://doi.org/10.1016/j.cub.2015.10.025

Sze, H., 1985. H+-Translocating ATPases: Advances Using Membrane Vesicles. Annu. Rev. Plant. Physiol. 36, 175–208. https://doi.org/10.1146/annurev.pp.36.060185.001135

Sze, H., Li, X., Palmgren, M.G., 1999. Energization of Plant Cell Membranes by H+-Pumping ATPases: Regulation and Biosynthesis. Plant Cell 11, 677–689. https://doi.org/10.1105/tpc.11.4.677

Tretyn, A., Wagner, G., Felle, H.H., 1991. Signal Transduction in Sinapis alba Root Hairs: Auxins as External Messengers. Journal of Plant Physiology 139, 187–193. https://doi.org/10.1016/S0176-1617(11)80606-1

Uchida, N., Takahashi, K., Iwasaki, R., Yamada, R., Yoshimura, M., Endo, T.A., Kimura, S., Zhang, H., Nomoto, M., Tada, Y., Kinoshita, T., Itami, K., Hagihara, S., Torii, K.U., 2018. Chemical hijacking of auxin signaling with an engineered auxin–TIR1 pair. Nat Chem Biol 14, 299–305. https://doi.org/10.1038/nchembio.2555

Ullrich, C.I., Novacky, A.J., 1990. Extra-and Intracellular pH and Membrane Potential Changes Induced by K+, Cl−, H2PO4−, and NO3− Uptake and Fusicoccin in Root Hairs of Limnobium stoloniferum. Plant Physiol. 94, 1561–1567. https://doi.org/10.1104/pp.94.4.1561

von Wangenheim, D., Hauschild, R., Fendrych, M., Barone, V., Benková, E., Friml, J., 2017. Live tracking of moving samples in confocal microscopy for vertically grown roots. eLife 6. https://doi.org/10.7554/eLife.26792

Wickham, H., 2007. Reshaping Data with the reshape Package. J. Stat. Soft. 21. https://doi.org/10.18637/jss.v021.i12

Yamada, R., Murai, K., Uchida, N., Takahashi, K., Iwasaki, R., Tada, Y., Kinoshita, T., Itami, K., Torii, K.U., Hagihara, S., 2018. A Super Strong Engineered Auxin-TIR1 Pair. Plant Cell Physiol 59, 1538–1544. https://doi.org/10.1093/pcp/pcy127

