## Supplemental 1 for "The AFB1 auxin receptor controls rapid auxin signaling and root growth through membrane depolarization in *Arabidopsis thaliana*"

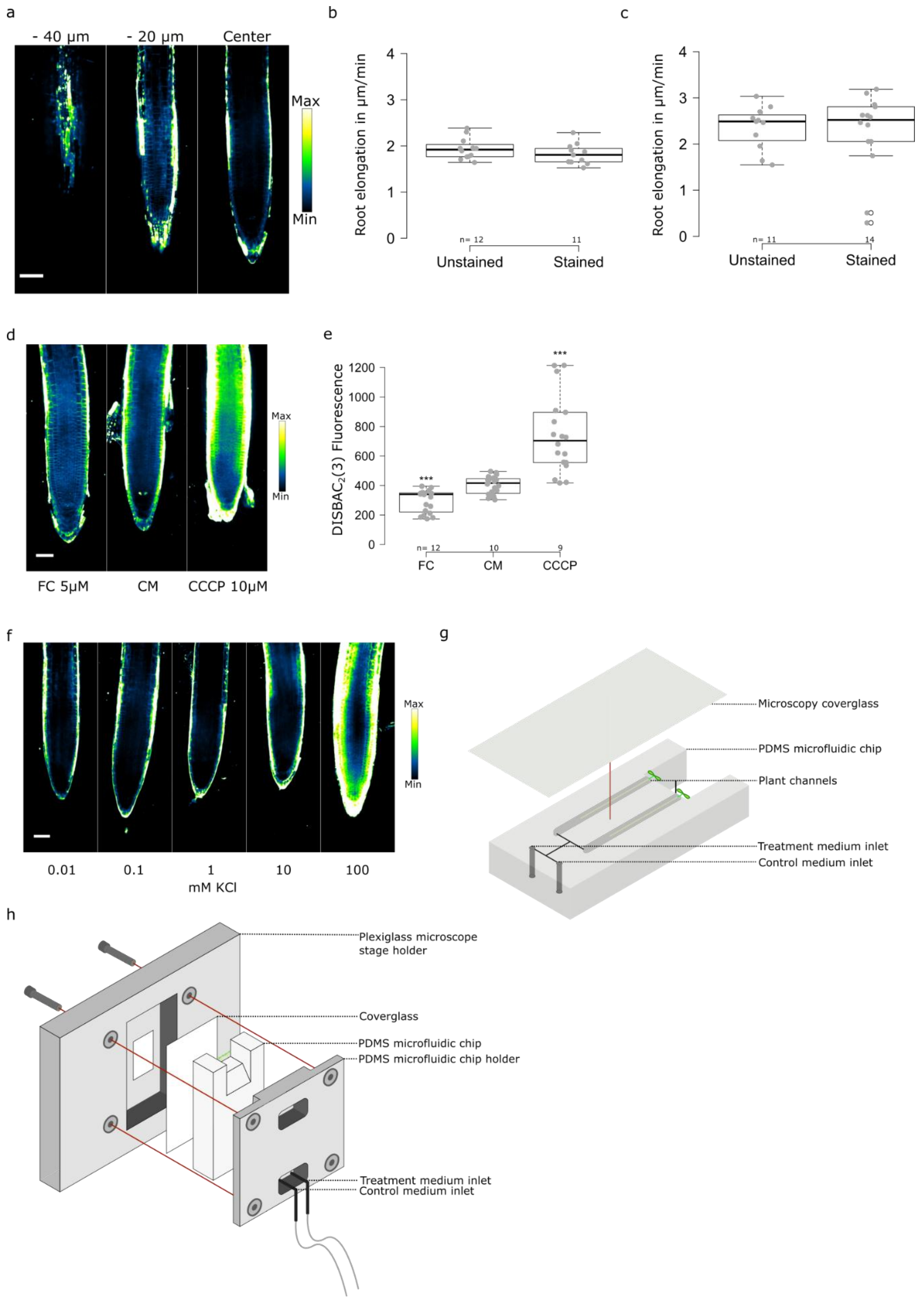


**Figure S1.** a) Z profile of DISBAC$_2$(3) fluorescence from the epidermis (-40µm) to the center of the root. b,c) Effect of DISBAC$_2$(3) staining (0µM and and 15µM) on primary root elongation (µm/minute) measured b) every 30 minutes over 10 hours grown on agar surface and b) measured every 5 minutes over 40 minutes and imaged using the spinning disk microscope. d,e) Effect of 5µM FC and 10µM CCCP compared to control (CM) on DISBAC$_2$(3) fluorescence after 20 minutes of treatment; d) representative images and e) fluorescence quantification. f) DISBAC$_2$(3) fluorescence in response to a KCl gradient (0.01 to 100mM by x10 increments). g,h) Schematic of the closable PDMS microfluidic chip. Representative microscopy images, fluorescence look up table indicated next to images, scale bars = 50µm. n= are indicated on figures. ***: p-value<0.0005.
