## Supplemental 2 for "The AFB1 auxin receptor controls rapid auxin signaling and root growth through membrane depolarization in *Arabidopsis thaliana*"

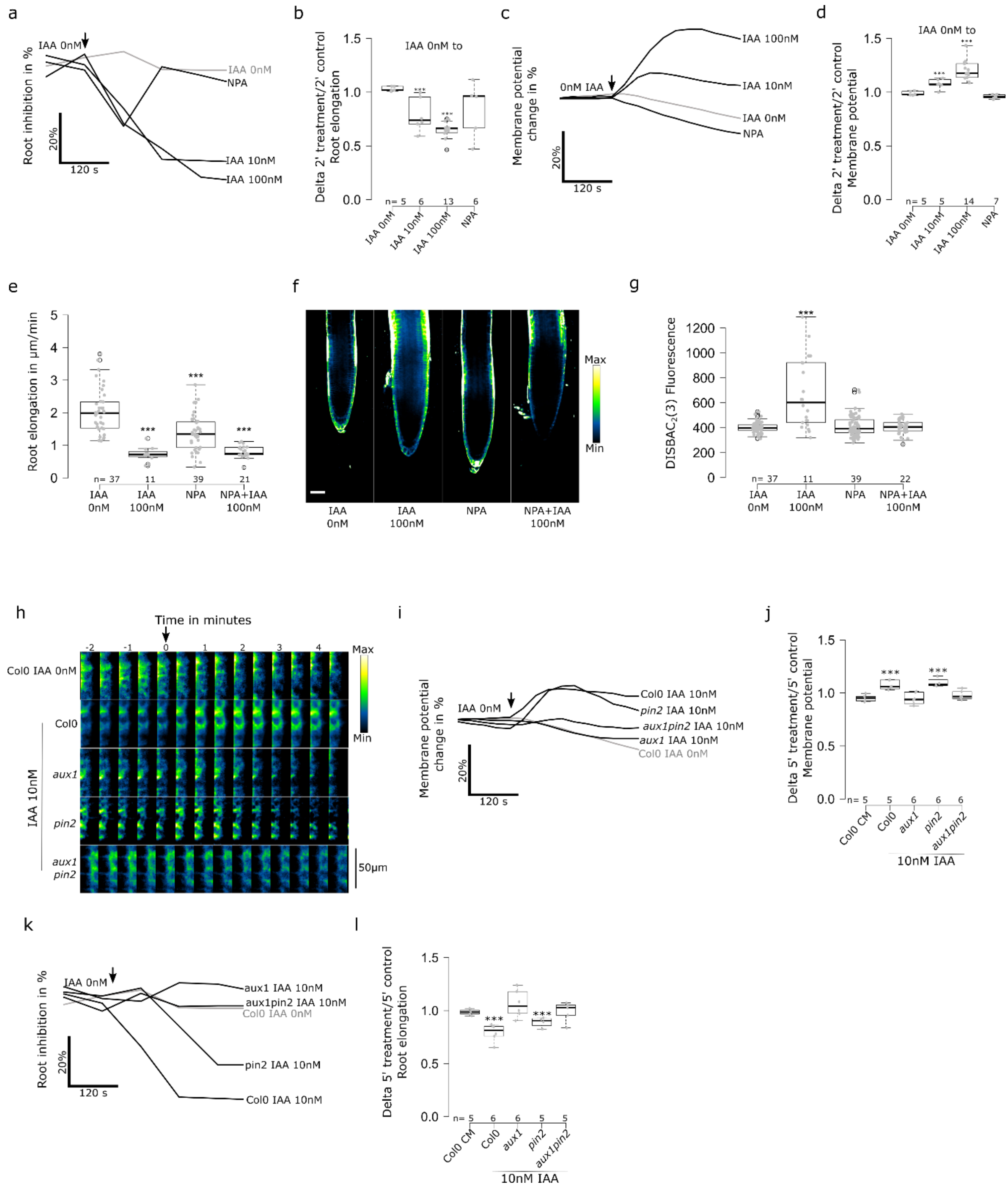

**Figure S2.** a-d) Effect of 20µM NPA on rapid auxin response. a) Root elongation over 5 minutes of treatment; average change (in %) over 5 minutes of treatment and b) delta of 2 minutes treatment over 2 minutes control media. c) Mean change of membrane potential (in %) over 5 minutes of treatment and d) delta of 2 minutes treatment over 2 minutes control media. e-g) Effects of NPA and IAA on steady state e),f) root tip DISBAC$_2$(3) fluorescence and g) primary root elongation (µm/minute). h-k) Effect of 100nM IAA on the rapid response of Col0, *aux1*, *pin2* and *aux1pin2.* h) mean change in membrane potential (in %) over 5 minutes of treatment and i) delta of 5 minutes treatment over 5 minutes control media. j) Root elongation over 10 minutes of treatment, mean change (in %) over 5 minutes of treatment and k) delta of 5 minutes treatment over 5 minutes control media. Steady state corresponds to the fluorescence of roots after 20 minutes of treatment in agar medium and root elongation measured over 40 minutes. Rapid response corresponds to roots treated in microfluidics and measured every 30 seconds. Application of treatments is indicated by a black arrow. For f and h, standard errors were not added to simplify reading. The selected microscopy pictures are representing the median fluorescence value. Fluorescence look up table indicated next to images, scale bar = 50µm. n= is indicated on figures. ***: p-value<0.0005.
