## Supplemental 3 for "The AFB1 auxin receptor controls rapid auxin signaling and root growth through membrane depolarization in *Arabidopsis thaliana*"

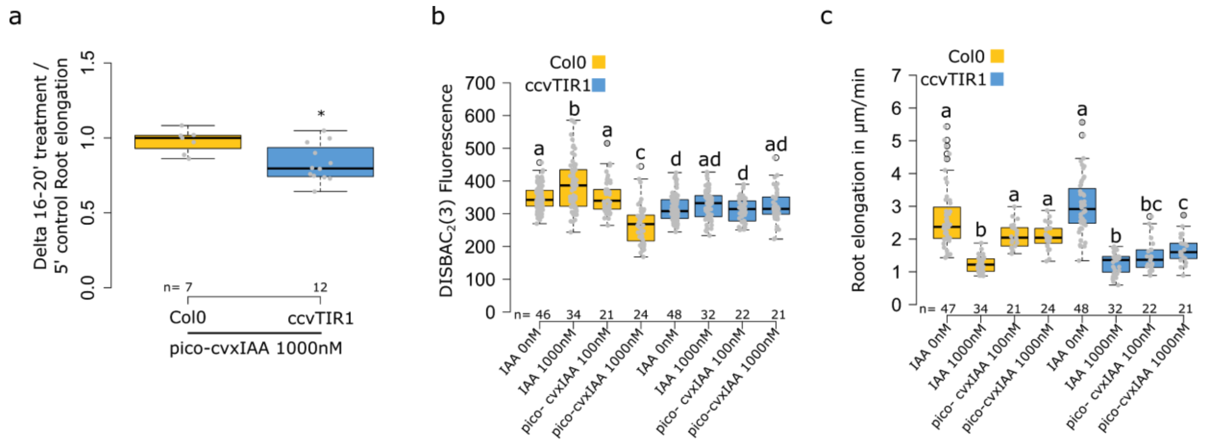


**Figure S3.** a) Effect of pico-cvxIAA on root elongation ratio of 16-20 minutes treatment over 5 minutes control media during microfluidics experiment. b,c) Effect of IAA and pico-cvxIAA on the steady state response of Col0 and ccvTIR1 b) root tip membrane potential and c) primary root elongation (µm/minute). Steady state corresponds to the fluorescence of roots after 20 minutes of treatment in agar medium and root elongation measured over 40 minutes. n= is indicated on figures. ***: p-value<0.0005, *: p-value<0.05.
