## Supplemental 4 for "The AFB1 auxin receptor controls rapid auxin signaling and root growth through membrane depolarization in *Arabidopsis thaliana*"

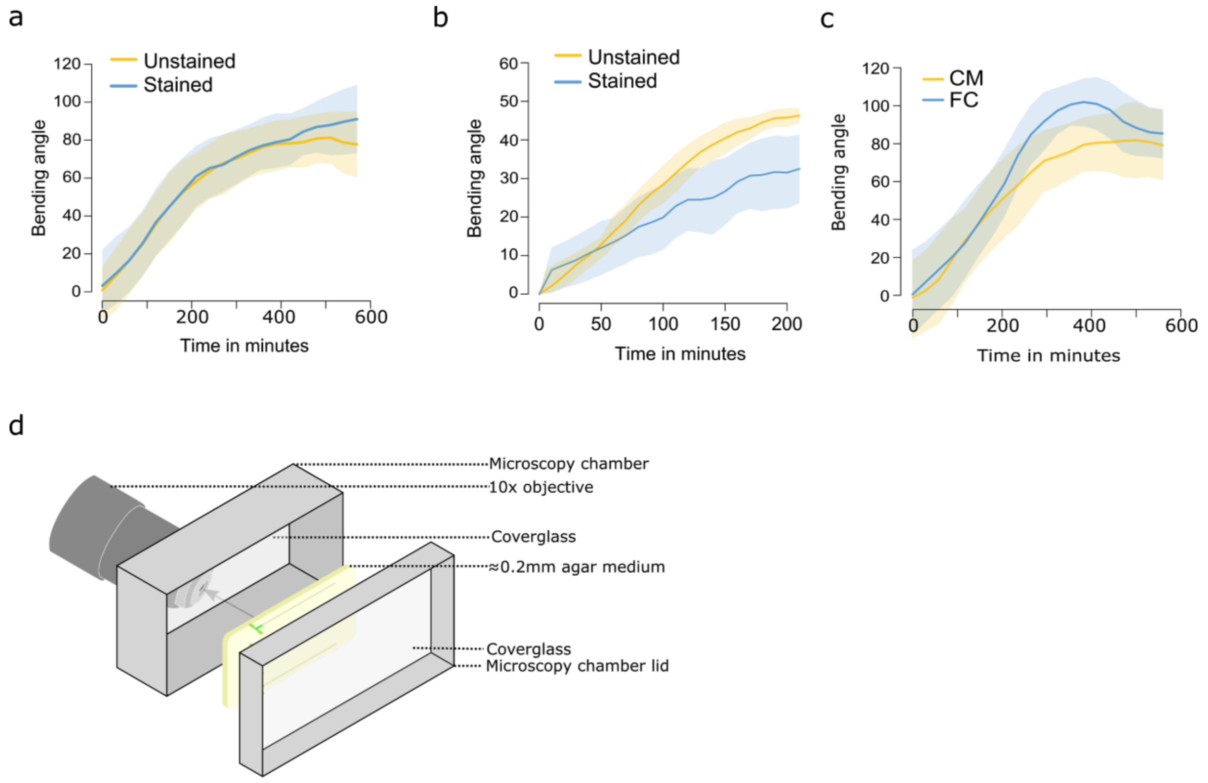


**Supplemental 4.** Effect of $DISBAC_2(3)$ staining (0µM and 15µM) on gravitropic root bending a) root tip angle measured every 30 minutes over 10 hours grown on agar surface and b) t measured every 5 minutes over 40 minutes grown in the imaging chamber on the spinning disk microscope. c) quantification of the effect of root tip angle after 90° gravistimulation of Col0 in control media (CM) or 5µM fusicoccin (FC). Images were taken every 30 minutes for 550 minutes. d) Schematic of the setup for high spatio-temporal resolution measurements of gravitropic bending with a vertical stage microscope.
