## Supplemental movie captions for "The AFB1 auxin receptor controls rapid auxin signaling and root growth through membrane depolarization in *Arabidopsis thaliana*"

**Supplemental movie 1.** Membrane potential of a Col0 root growing in microfluidic channels and treated with 100nM IAA. Scale bar = 50µm. Treatment starts at time 0. Time is minutes:seconds.

**Supplemental movie 2.** Col0 root tip angle measurements over 30 minutes after 90° gravistimulation. Angle value between (in degrees) the orange and blue line is indicated in the top left corner. Time in minutes is indicated in the bottom left corner.

**Supplemental movie 3.** *afb1-3* root tip angle measurements over 30 minutes after 90° gravistimulation. Angle value (in degrees) between the orange and blue line is indicated in the top left corner. Time in minutes is indicated in the bottom left corner.
